# Macrophages maintain signaling fidelity in response to ligand mixtures

**DOI:** 10.1101/2025.03.27.645368

**Authors:** Xiaolu Guo, Supriya Sen, Julian Gonzalez, Alexander Hoffmann

## Abstract

As immune sentinel cells, macrophages are required to respond specifically to diverse immune threats and initiate appropriate immune responses. This stimulus-response specificity (SRS) is in part encoded in the signaling dynamics of the NFκB transcription factor. While experimental stimulus-response studies have typically focused on single defined ligands, in physiological contexts cells are exposed to multi-ligand mixtures. It remains unclear how macrophages combine multi-ligand information and whether they are able to maintain SRS in such complex exposure conditions. Here, we leveraged an established mathematical model that captures the heterogeneous single-cell NFκB responses of macrophage populations to extend experimental studies with systematic simulations of complex mixtures containing up to five ligands. Live-cell microscopy experiments for some conditions validated model predictions but revealed a discrepancy when TLR3 and TLR9 are stimulated. Refining the model suggested that the observed but unexpected ligand antagonism arises from a limited capacity for endosomal transport which is required for responses to CpG and pIC. With the updated model, we systematically analyzed SRS across all combinatorial-ligand conditions and employed three ways of quantifying SRS involving trajectory decomposition into informative trajectory features or machine learning. Our findings show that macrophages most effectively distinguish single-ligand stimuli, and distinguishability declines as more ligands are combined. However, even in complex combinatorial conditions, macrophages still maintain statistically significant distinguishability. These results indicate a robustness of innate immune response specificity: even in the context of complex exposure conditions, the NFκB temporal signaling code of macrophages can still classify immune threats to direct an appropriate response.

**Significance (≤120):** Macrophages sense diverse pathogens within complex environments and respond appropriately. Experimental studies have found that the NFκB pathway responds with stimulus-specific dynamics when macrophages are exposed to single ligand stimuli. However, it remains unclear complex contexts might erode this stimulus-specificity. Here we systematically studies NFκB responses using a mathematical model that provides simulations of the heterogeneous population of single cell responses. We show that although the model is parameterized to single ligand data it can predict the responses to multi-ligand mixtures. Indeed, model validation uncovered signaling antagonism between two ligands and the underlying mechanism. Importantly, we found that NFκB signaling dynamics distinguish ligands within multi-ligand mixtures indicating a robustness of the NFκB temporal code that was not previously appreciated.

## INTRODUCTION

Functioning as sentinel immune cells, macrophages detect diverse immune threats by recognizing pathogen-associated molecular patterns (PAMPs), damage-associated molecular patterns (DAMPs) and cytokines (1, 2). In response, they initiate immune defenses tailored to the specific threat, including cell-intrinsic defenses, localized immune responses, or systemic immune activation (3, 4). The regulatory machinery governing these stimulus-specific functions converges on a few signaling pathways, such as IRF, AP1, NFκB, and p38 (5). Among these, NFκB is a key transcription factor that responds to all pathogens in macrophages and regulates immune gene expression. The temporal dynamics of NFκB encode environmental cues that characterize immune threats (6–9). These dynamic features can be quantified by signaling codons, which provide information about specific stimuli and direct distinct immune responses. Dysfunction in NFκB stimulus-response specificity has been associated with diseases such as Sjögren’s syndrome (8).

While experimental studies of stimulus-response have typically focused on single, chemically defined ligands (8–14), physiological stimulation from pathogens and injury often involves multiple chemical ligands. For example, BCG vaccines can activate immune cells via TLR2 and TLR4 (15). Additionally, organisms may encounter multiple immune threats simultaneously—such as concurrent viral and bacterial infections, in conjunction with cytokines —resulting in complex mixtures of chemical ligands. A few studies have explored representative combinations of such stimuli, such as bacterial infections involving co-stimulation of TLR2 and TLR4 (6) or bacterial infections along with host-responsive cytokines involving TLR4 and TNFR (16). However, due to the vast exploration space, there is a lack of systematic experimental investigation into stimulus-response specificity (SRS) in the context of ligand mixtures across all possible combinations. This challenge could be addressed through computational simulations so long as a reliable mathematical model is available that has been parameterized to account for the heterogeneous single-cell stimulus response trajectories found in the population of macrophages.

There is an extensive mathematical modeling literature of the NFκB regulatory mechanisms. The early models captured bulk-average NFκB dynamics in different IκB knockout genotypes (17), and explored the mechanisms that could account for single cell trajectories (18–20, 8). Significant modeling efforts have been made to account for the cell to cell heterogeneity through parameter selection (10, 13, 21–24). These mechanistic models, which integrate extensive experimental data of protein interactions, translocation, or catalysis, have provided confidence in the accuracy of the underlying biochemical reaction network. Building on this foundation, we have recently developed model simulations of a population of heterogeneous single-cell NFκB signaling trajectories. A quantitative assessment of this model demonstrates that it successfully captures the experimental stimulus-response specificity (SRS) for single-ligand stimuli while accounting for cellular heterogeneity within the population (25).

Here we have leveraged these advances to predict NFκB responses to combinatorial ligands. A key challenge which is that, for each parameterized virtual single cell, only one receptor module (comprising the regulatory network that is receptor proximal) and the shared IKK-IκB-NFκB core module can be inferred from single-ligand experiments. Information about other receptor modules remains unknown for that same cell. To address this, we developed a workflow that infers missing receptor module information from other virtual cells based on similarity in core module parameters (Figure 1A) (25). In this work, we employed this model to simulate combinatorial-ligand responses to explore SRS in context of ligand mixture. With model simulation and then experimental validation, we found that the model simulations are reliable for many, but not for all conditions, providing opportunities to improve our understanding of regulatory mechanisms, improve the model iteratively, and address how well the NFκB temporal signaling code distinguishes complex mixtures of ligands.

**Figure 1.**
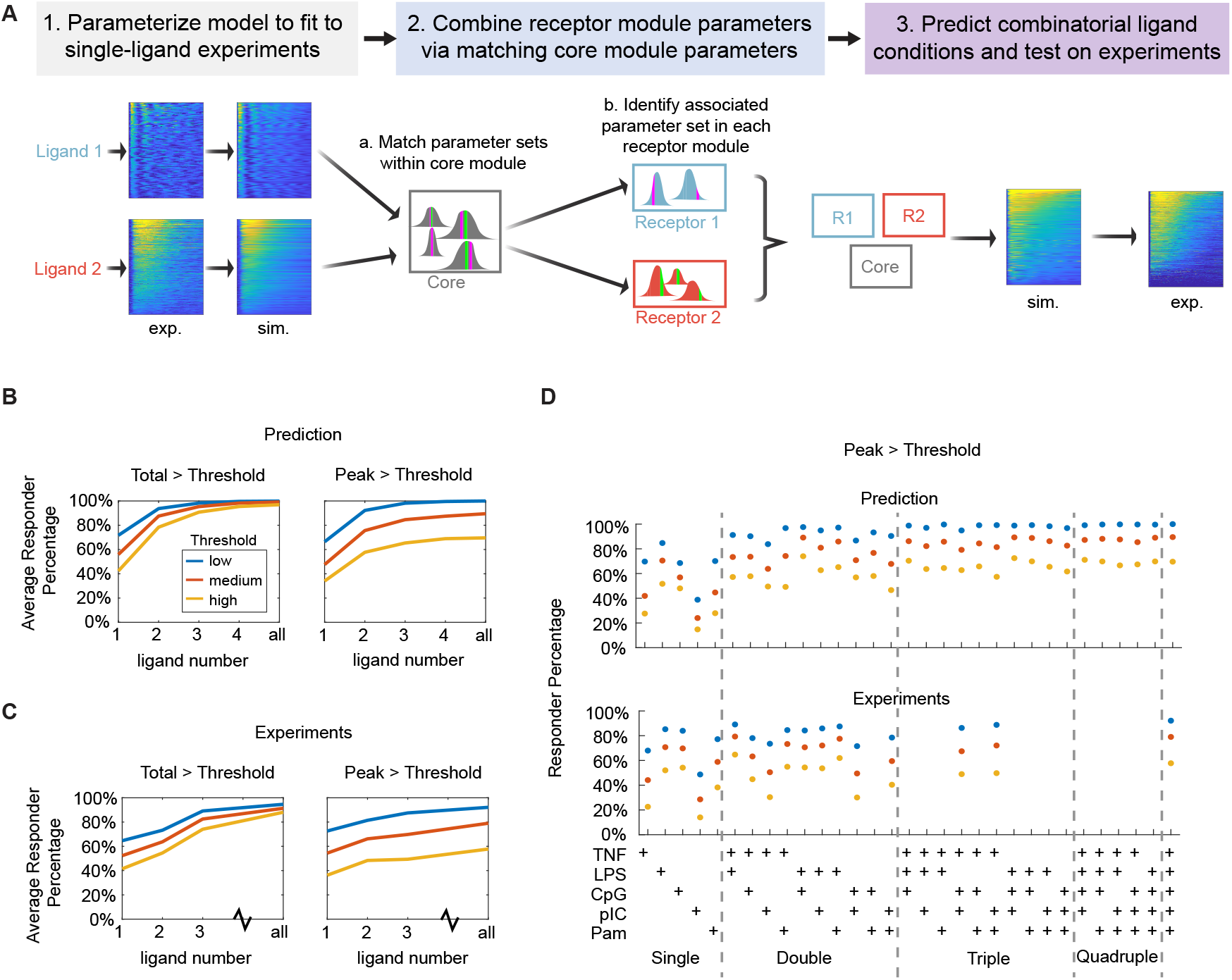
Extending a single-ligand-trained mathematical model to predict responses to ligand mixtures. A. Schematic of the workflow: A mathematical model is parameterized to account for heterogenous NFκB signaling dynamics in response to five single-ligand stimulation conditions. This involves parameters within the regulatory networks of the ligand-specific receptor module and the shared IKK-IκB-NFκB core module. By matching similar core module parameter sets, specific parameter sets for the two receptor modules are combined, thereby enabling simulations of NFκB responses to ligand mixtures. B. Model-predicted average responder fraction for all single, double, triple, quadruple or the five ligand stimulations. Responder fractions are defined by low, medium or high thresholds applied to either Total (left) or Peak (right). C. Experimentally determined average responder fraction for the single, double, triple, quadruple or all ligands as determined in panel D. D. Responder fractions defined by a low, medium, or high threshold applied to Peak activity for each indicated ligand and ligand combination. These data define the averages shown in B and C.

## RESULTS

### Predicting NFκB responses to combinatorial ligand stimuli

To study NFκB response dynamics in response to stimulation with combinatorial ligands, we employed a well-parameterized single-cell mechanistic model of the NFκB signaling network (25). This mechanistic model was trained on single-ligand response datasets, capturing the single-ligand stimulus-response specificity of the population of macrophages while accounting for cellular heterogeneity. The five ligands include the cytokine Tumor Necrosis Factor (TNF); TLR2 ligand Pam3CysSerLys4 (Pam), TLR9 ligand Cytosine-phosphate-Guanine (CpG), and TLR4 ligand Lipopolysaccharide (LPS), which are bacterial PAMPs; and TLR3 ligand Polyinosinic:polycytidylic acid (pIC), which is a viral PAMP.

We employed a workflow to predict combinatorial ligand stimulation from single-ligand stimulated data. In this workflow, we first determined parameter distributions for the core module and the cognate receptor module that provided the best fit for the single ligand experimental data (Figure 1A, Step 1). Next, we combined the five single-stimulus models into multi-stimulus models by matching cells with similar core module parameters. Since the IκB-NFκB core module is shared across all stimuli, we matched core modules across different stimulus conditions by minimizing parameter Euclidean distance metrics between conditions (Figure 1A, Step 2). Specifically, for a given cell *i* responding to ligand *l* (*i* ∈ *S*_*l*_), we identified a corresponding cell *j* responding to ligand *k* (*j* ∈ *S*_*k*_) through the following optimization:

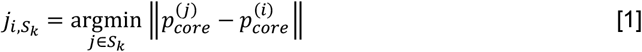

where 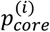 represents the core parameter vector of cell *i*. The parameters of cell *i* in the receptor module associated with ligand *k* (hereafter referred to as receptor module *k*) were then inferred as:

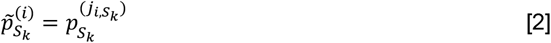

where 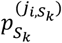 represents the parameters of receptor module *k* for cell 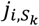, and 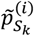 represents the extended receptor module *k* parameters of cell *i*.

Using this approach, we simulated NFκB signaling responses under various conditions, including five single-ligand stimuli (TNF, Pam, CpG, LPS, and pIC), ten double-ligand combinations, ten triple-ligand combinations, five quadruple-ligand combinations, and one quintuple-ligand condition—a total of 31 conditions. The model was trained exclusively on single-ligand data (red box, Figure S1A) and used to predict combinatorial responses (blue box, Figure S1A).

To quantify NFκB dynamics, we decomposed trajectories into six signaling codons (Speed, Peak, Duration, Total (integral), EvL (early loaded vs late activity), and Osc (oscillation content)), which were identified as the informative features of NFκB trajectories (Figure S1B) (8). Heatmaps and signaling codon distributions revealed that combinatorial-ligand conditions were generally predicted to exhibit stronger responses (Total) compared to single-ligand conditions.

### Responder ratio increase with ligand numbers for combinatorial ligand stimuli

To test our model predictions, we conducted experimental studies (Figure S1C). Both simulations and experimental data indicate the presence of non-responding cells under single-ligand conditions (Figure S1A). With this approach we aimed to address the question: do non-responders observed experimentally arise from general cellular ill-health, or are they due to cellular heterogeneity in the ligand-specific receptor modules?

To address this, we defined the responder ratio as the fraction of cells with Peak or Total NFκB signaling codons exceeding predefined thresholds. The model predicted that the responder fraction generally increases for combinatorial conditions compared to single-ligand stimuli (Figure 1B). When applying thresholds on Total activity, the average responder ratio reaches a saturation point of over 90% that is largely independent of the threshold applied. When applying thresholds on Peak activity, the average responder fraction also increases but the plateau is sensitively dependent on the threshold. We analyzed the experimental data analogously (Figure 1C) and found remarkable concordance; the responder fraction increases with the number of ligands used, reaching a plateau that is either independent of threshold (Total) or dependent on threshold (Peak). This analysis provides confidence in the validity of the model as well as the experimental workflow. It suggests that signaling from multiple ligands generally combine in an integrative manner, and that non-responder cells are likely a result of cellular heterogeneity in receptor modules rather than the core module.

Examining the individual stimulus conditions in detail, we found that the responder fraction varies substantially between ligands (e.g. lowest for pIC and highest for LPS) as well as between ligand mixtures (Figure 1D). For the available experimental data, we found good agreement with the model predictions. Some discrepancies between simulations and experiments were observed, particularly in cases of non-integrative responses, where the combinatorial-ligand response is lower than at least one of the corresponding single-ligand responses. For instance, the model predicted that CpG-pIC would exhibit a comparable response to pIC-Pam and a higher response than CpG or pIC alone (Figure 1D, top panel). However, experimental results showed a non-integrative response for the CpG-pIC pair, with lower activation than CpG alone and significantly lower than pIC-Pam (Figure 1D, bottom panel).

### Limited endosomal transport can result in ligand antagonism

To investigate the discrepancy in more detail, we examined the simulated and experimentally measured trajectories of TNF+LPS, pIC+LPS, and pIC+CpG dual-ligand simulations and corresponding single-ligand stimulations (Figure 2A). Experiments showed good concordance for LPS+pIC and TNF+LPS, but not CpG+pIC, which showed much lower responses than predicted (Figure 2B). By comparing signaling codon distributions calculated for simulated and experimental trajectories (Figure S2) we similarly found good general agreement, although some discrepancies, such as for the CpG-pIC pair, were evident. Specifically, we found that LPS+pIC, and TNF+LPS responses to be integrative as predicted (Figure 2C). However, the CpG+pIC stimulated response was non-integrative: both Duration and Total were lower for CpG+pIC than for CpG alone, in contrast to model predictions.

**Figure 2.**
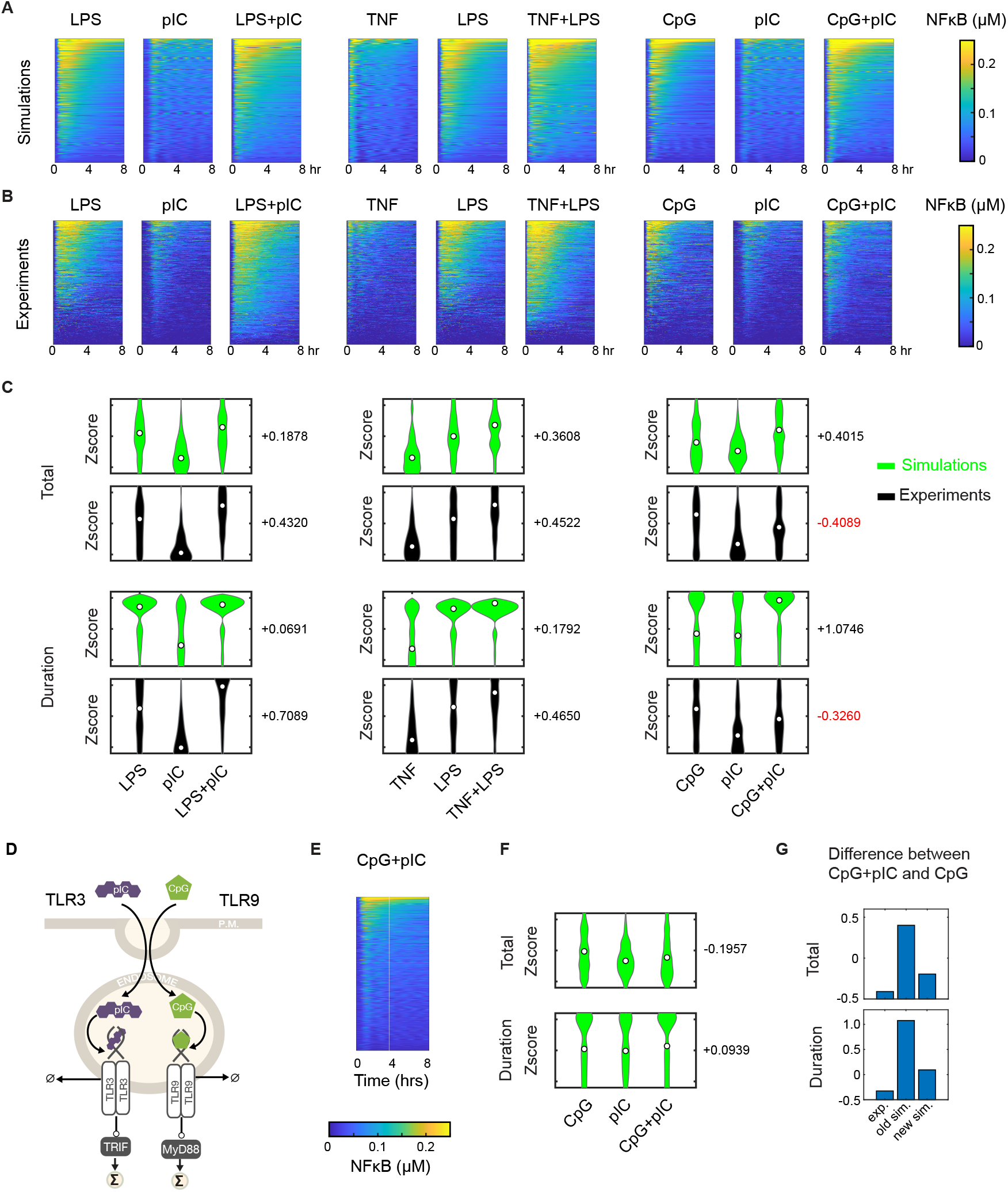
Iterative modeling and experimentation of combinatorial ligand stimulation. A. Heatmaps of model-predicted NFκB signaling trajectories in macrophages responding to combinatorial ligand stimulation: LPS+pIC, TNF+LPS, and CpG+pIC. Trajectories were ordered by Total and color intensity indicates nuclear NFκB abundance in accordance with the color bar. B. Heatmaps of experimental data for the conditions shown in (B). C. Violin plots comparing signaling codon distributions of Total (top) and Duration (bottom) from single and combinatorial ligand stimulation, as predicted by the model (green) and determined by experiments (black). The hollow circle in each plot indicates the distribution median. D. Schematic of the regulatory network in which CpG and pIC use the same endosomal transport machinery. The potential saturation of endosomal transport may lead to functional antagonism between the two ligands. E. Heatmap of predicted NFκB signaling trajectories in macrophages upon combinatorial stimulation with CpG and pIC under transport saturation conditions. F. Violin plots of the signaling codon distributions of Total (top) and Duration (bottom) as predicted by the refined model for single and combinatorial ligand stimulation. G. Bar graph of the mean value difference in the Total (left) and Duration (right) signaling codon distributions shown in panel C for oirginal model and in panel E for refined model.

To address the model’s shortcoming, we noted that both stimuli are sensed by receptors located in the endosomal compartment. The endosomal transport machinery may have limited capacity (26, 27) but the model represented these as independent mass action reactions. We reformulated the model of CpG and pIC receptor modules with a Hill equation that describes the limited capacity of the common endosomal trafficking process in their signaling pathways (Figure 2D, S3), such that only one ligand may saturate it.

The updated model introduced a negative correlation between endosomal CpG-receptor and pIC-receptor complexes due to competition for limited endosomal transport (Figure S3) and the fact that pIC-TRIF signaling is generally weaker than CpG-MyD88 signaling. Simulations with the revised model accurately reflected the non-integrative CpG+pIC response (Figure 2E), where Duration and Total activity for the combined stimuli were lower than for CpG alone (Figure 2F-G). In sum, our results suggest that saturation of endosomal transport may lead to non-integrative signaling responses for stimulus ligands that are sensed by endosomal receptors.

### Stimulus-response specificity for combinatorial ligand stimuli

We employed the updated model to simulate NFκB signaling dynamics and decomposed the trajectories into signaling codons (Figure S4A-B), to enable investigations of the stimulus response specificity (SRS). We first calculated the Wasserstein distance between signaling codons in different stimulus conditions, as this metric has been used to quantify the similarity and specificity across conditions (25). This confirmed a high distinguishability for single-ligand stimuli but revealed that the distinguishability diminishes with the number of combinatorial ligands that make up the stimulus condition (Figure 3A).

**Figure 3.**
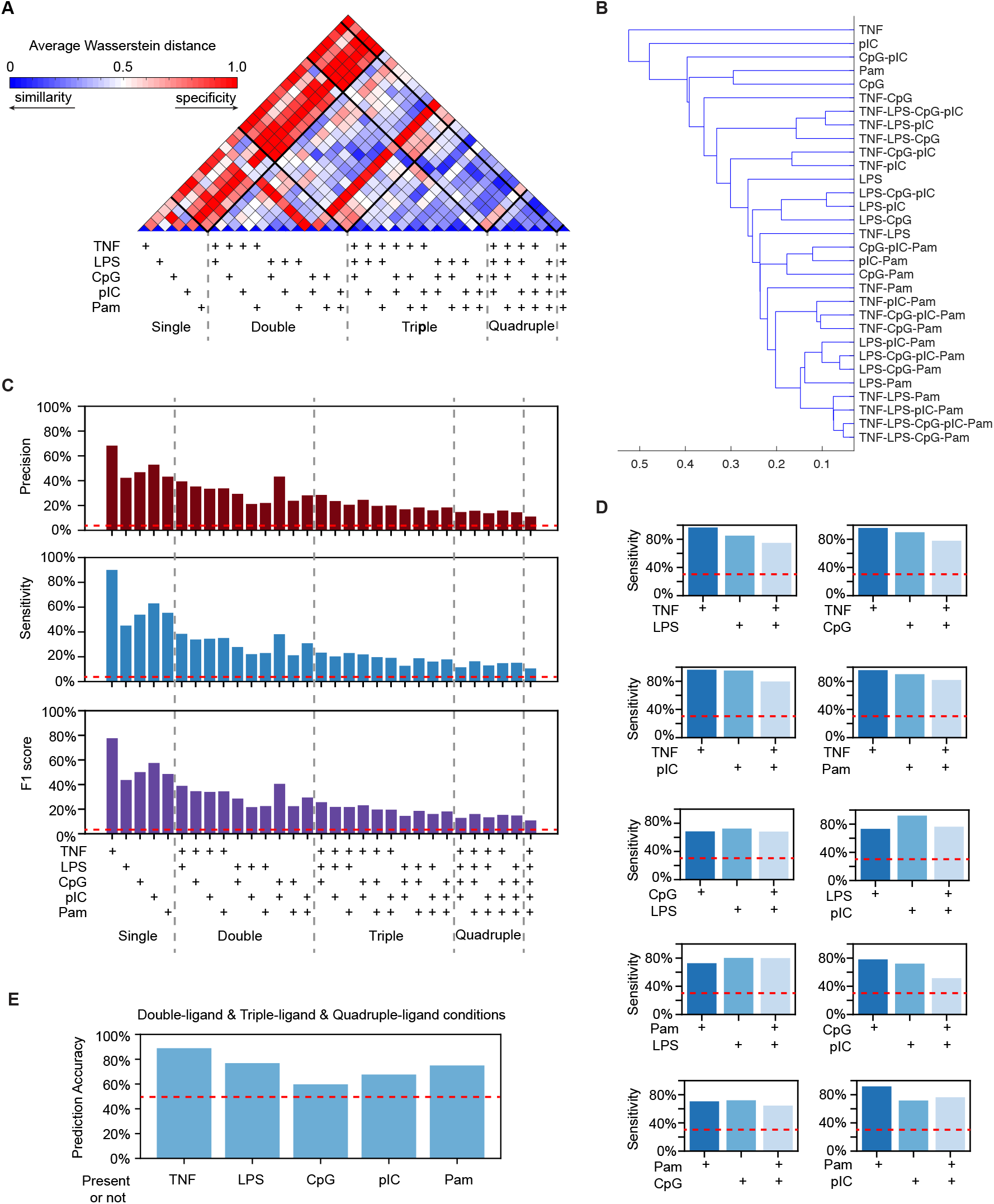
Distinguishability of single-ligand stimuli and combinatorial ligands stimuli. A. Heatmap of average Wasserstein distance between the distributions of 6 NFκB signaling codons for each of 31 conditions. Conditions are identified on the diagonal axis. Color intensity indicates average Wasserstein distance in accordance with the color bar. B. Hierarchical clustering of the 31 conditions *via* single-linkage clustering of the average Wasserstein distance metric. C. Bar plots of Precision, Sensitivity, and F1 score for a Random Forest classifier trained on all 31 conditions using NFκB response signaling codons as input to predict the stimuli. 67% of the data was used for training and 33% for testing. D. Bar plots of Sensitivity for Random Forest classifiers trained on all 3 conditions (each single ligand and the pair). Methodology as in panel C. E. Bar plots depicting the prediction accuracy of classifiers trained on TNF-present vs. TNF-absent, LPS-present vs. LPS-absent, CpG-present vs. CpG-absent, pIC-present vs. pIC-absent, and Pam-present vs. Pam-absent groups generated from all double-ligand, triple-ligand, and quadruple-ligand conditions. Methodology as in C.

To rank the most distinguishable stimuli among ligand mixtures, we applied hierarchical clustering to all 31 conditions based on their Wasserstein distances (Figure 3B). TNF was the most distinguishable ligand in context of all possible ligand mixtures (with hierarchical height ≈ 0.5 to the nearest cluster), followed by the viral PAMP pIC (hierarchical height ≈ 0.49 to the nearest cluster). Bacterial PAMPs (LPS, CpG, and Pam) are less distinguishable compared to TNF or pIC, but still distinguishable from other mixtures of ligands (average hierarchical height ≈ 0.3 to the corresponding nearest cluster). Overall, single-ligand conditions were more specific than combinatorial-ligand conditions.

To further examine SRS in ligand mixtures, we trained a random forest classifier and quantified the specificity for each condition via different prediction accuracy metrics. Machine learning classifiers have been widely used to assess the distinguishability and confusion in signaling dynamics (8, 9). Analysis of the classifier’s precision and sensitivity matrix showed that true positive rates (diagonal elements, quantifying specificity) were substantially higher than misclassification rates (off-diagonal elements, quantifying the confusion between conditions) (Figure S5A-B). When plotting only the true positives, single-ligand conditions demonstrated the highest sensitivity, precision, and F1 scores, while distinguishability declined with increasing ligand numbers (Figure 3C, S5A-B). Notably, the TNF-stimulated condition achieved the highest distinguishability with approximately 70% precision, 90% sensitivity, and an 80% F1 score, whereas the five-ligand combination condition exhibited the lowest distinguishability (≈10% for sensitivity, precision, and F1 score), though still far above the random guess baseline of 3.2% (red dashed line in Figure 3C). We also applied a Long Short-Term Memory (LSTM) machine learning model to the entire time-series trajectories (28), and found similar results, confirming signaling codons capture all the information present in the newly simulated data (Figure S6).

We then used machine learning classifiers to investigate whether dual-ligand conditions show confusion with the corresponding single-ligand conditions. For each ligand pair, a model was trained to distinguish among three conditions: ligand A alone, ligand B alone, and the combined stimulus (ligand A+B). While dual-ligand stimuli were less distinguishable than their respective single-ligand counterparts, they remained well differentiated, with sensitivities ranging from 50% to 80% (Figure 3D) and precisions from 60% to 90% (Figure S7), significantly outperforming the 33% expected from random guessing (red dashed line in Figure 3D and S7). Among all pairs, TNF-alone condition exhibited the highest specificity in the TNF-related pairs, achieving over 95% sensitivity and more than 90% precision. TNF stimulation remained the most distinguishable, even from TNF-involved dual-ligand combinations.

Ultimately, the cell needs to determine whether specific immune stimuli are present. To test this, we grouped all double-ligand, triple-ligand, and quadruple-ligand conditions (in total 25 conditions) into two categories: one with 14 conditions where TNF is present and another with 11 conditions where TNF is absent. We then trained machine learning classifiers to predict whether a cell was exposed to TNF based on its NFκB signaling codon. The results showed that classifiers trained on the TNF-present vs. TNF-absent groups achieved 90% prediction accuracy, significantly higher than the random-guess baseline of 50% (Figure 3E). We repeated this analysis for other ligands, including pIC, LPS, CpG, Pam. The results demonstrated that cells could reliably determine the presence of LPS with 76% accuracy, Pam with 75% accuracy, pIC with 65% accuracy, and CpG with 59% accuracy in ligand mixtures (Figure 3E).

## DISCUSSION

We report here on using a mathematical model to extend the insights gained from an experimental model and learn additional information about macrophage stimulus-response functions. We found that NFκB, the primary transcription factor responding to inflammatory and innate immune stimuli and known to do so stimulus-specifically, maintains a degree of stimulus-specificity in its responses when cells are exposed to complex ligand mixtures. Complex ligand mixtures are thought to be more physiologically relevant than stimuli consisting of a single defined ligand. To enable these studies, we first iteratively refined the model with experimental data to identify an unexpected molecular mechanism. Specifically, we found that a limited endosomal transport capacity may lead to antagonism between ligands whose receptors are in the endosome. This signaling interaction in turn enables superior stimulus-response specificity of the respective stimulus-pair.

Mechanistic models have two advantages over statistical models which were illustrated in this work: by leveraging knowledge, they allow us to 1) extend the predictions to outside the training range and 2) understand mechanism. The NFκB signaling model is based on decades of experimental study by numerous laboratories (17, 18, 29, 19, 10, 20, 8). These prior studies focused on single ligands, but here we focused on combinations which represent stimulus conditions that are outside the model’s training range, and signaling outcomes of ligand pairs cannot be interpolated from single ligand responses. Further, the interpretability of mechanistic models allowed us to gain new biological insights when model predictions deviated from experimental results. We found that endosomal transport, which is a requirement for some ligands as receptors are located there (30, 31), can be a bottleneck, setting up the possibility for ligand antagonism. Other possibilities are shared signaling proteins such as MyD88 (23) or TRIF, or other shared resources such as core machineries controlling transcription, translation, or degradation.

We characterized the stimulus-response specificity (SRS) of NFκB signaling with several approaches. By decomposing the trajectories into the previously identified informative trajectory features termed signaling codons, we could calculate Wasserstein distances and train and evaluate a random forest machine learning model. Further, we applied a Long Short-Term Memory (LSTM) machine learning model in case additional information is present in the newly simulated data that is not represented by the previously identified signaling codons. The results were consistent: NFκB signaling dynamics provide for a degree of stimulus distinction even when those stimuli are complex ligand mixtures. However, the degree of SRS for ligand mixtures is lower than for the individual ligand and doses which were chosen in prior studies because they provide a high degree of SRS (8). The diminished SRS for ligand mixtures may also be rationalized by the fact that signaling is largely integrative and combines in a monotonic manner. There are exceptions, such as the antagonism identified for the CpG-pIC pair. Given there are likely other mechanisms that may lead to non-monotonicity that would be revealed in further model refinements, the SRS calculations presented here are likely underestimates. Our results therefore imply the robustness of innate immune response specificity that is maintained even when the immune threat occurs in the context of complex stimulus conditions.

Furthermore, innate immune responses do not solely rely on NFκB but also involve the critical functions of AP1, p38, and the IRF3-ISGF3 axis. The additional pathways are likely activated in a coordinated manner and provide additional information (5). In a previous study p38 did not add additional information to NFκB in single ligands, however these ligands had been selected to maximize the SRS of the NFκB trajectories (32), thus it is possible that p38 plays a more important role in response to other ligands or ligand mixtures. That may also be the case for the IRF3-ISGF3 axis, which only responds to a subset of ligands. The limitations of the present work should motivate future studies that expand the live-cell imaging and mechanistic modeling work to all pathways activated by innate immune stimuli and then reassess the SRS of complex ligand mixtures (6, 16), altered dosing (33), and potentially temporal phasing of ligands (34, 35). The present work not only provides motivation for such studies but a blueprint for analytical strategies and the role of mathematical models in extending the capabilities of experimental studies.

## Materials and Methods

### Mathematical model of NFκB dynamics and single-cell parameter distributions

A well-established ODE model of NFκB signaling (8, 25) was used to investigate NFκB dynamics in response to different stimuli. Detailed model structure is given in Supplementary dataset 1. Single-cell parameters enabled simulations that aligned with experimental data by capturing stimulus specificity and cell-to-cell heterogeneity of stimulus responses (25). The parameter values are given in Supplementary dataset 2 and served as the sampling distribution for model predictions.

### Predicting responses to ligand mixtures

For predicting responses to combinatorial ligand stimuli, receptor module parameter sets were matched based on core module parameter sets using equations [1–2]. The workflow of combinatorial ligand stimulation was adapted from (25) and is described in SI Appendix. For each condition, 1,000 cells were simulated using MATLAB’s ode15s solver. Simulations included an initial phase to establish steady-state, followed by stimulation. The resulting trajectories were visualized and analyzed in MATLAB.

### Experimental data generation

Myeloid precursor cells derived from the RelA-mVenus mouse strain (8) and transduced with HoxB4 (36) were cultured into macrophages using the standard bone marrow-derived macrophage (BMDM) culture method (8). The resulting macrophages were then stimulated with specified concentrations of LPS, TNF, CpG, pIC, and their combinations, without replacing the media. A live cell microscopy workflow allowed the measurement of nuclear NFκB activity at single-cell resolution, capturing images every 5 minutes over an 8-hour period. Fluorescence intensity was normalized to image background levels and baseline-corrected using an automated image analysis workflow, MACKtrack (https://github.com/Adewunmi91/MACKtrack).

### Signaling Codon decomposition and Responder fractions

We decomposed NFκB trajectories into six NFκB signaling codons—Speed, Peak, Duration, Total, EvL, and Osc—that encode stimulus information (8). The quantification was described (25) (see SI Appendix for details). Responder fractions were determined by imposing specified thresholds on Peak and Total.

### Model Refinement for CpG and pIC

To describe the use of a shared transport pathway by CpG and pIC we used Hill kinetics that may account for saturation. This is account for by multiplying the endosomal transportation rate by the inhibition Hill formula:

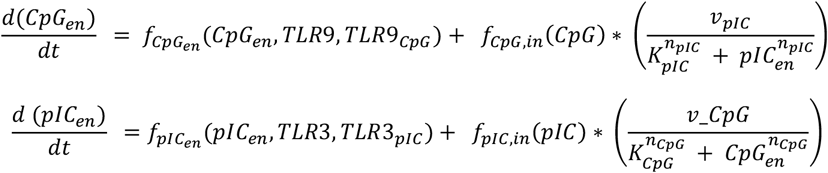

More details are described in SI Appendix.

### Stimulus-response specificity (SRS) assessment

To analyze stimulus-response specificity (SRS) of NFκB dynamics across different stimulations, we computed the distributions of signaling codons from simulated datasets using the updated CpG-pIC competition model. Wasserstein-2 distances were computed between codon distributions across different conditions, and the average distance across six signaling codons was used to quantify stimulus-response specificity. Using the simulated data, A random forest classifier was utilized to predict condition identity based on signaling codons, while LSTM classifiers were used to classify conditions based on NFκB dynamic trajectories. LSTM classifiers was used to predict conditions based on NFκB dynamics trajectories. More details are described in SI Appendix.

## Data availability

All experimental and simulation data are available at Mendeley Data (DOI:10.17632/bv957×6frk.1).

## Code availability

All codes are available at GitHub (https://github.com/Xiaolu-Guo/Combinatorial_ligand_NFkB).

## Acknowledgements

We thank Sarina Low and Allison Schiffman for critical reading the manuscript, and the Microscopy & Modeling floor meeting for suggestions. The work was supported by NIH grant R01AI173214 to AH.

