## Supplementary Information for "Macrophages maintain signaling fidelity in response to ligand mixtures"

### Supplementary Information: *Macrophages maintain signaling fidelity in response to ligand mixtures*

### Supplementary Figures

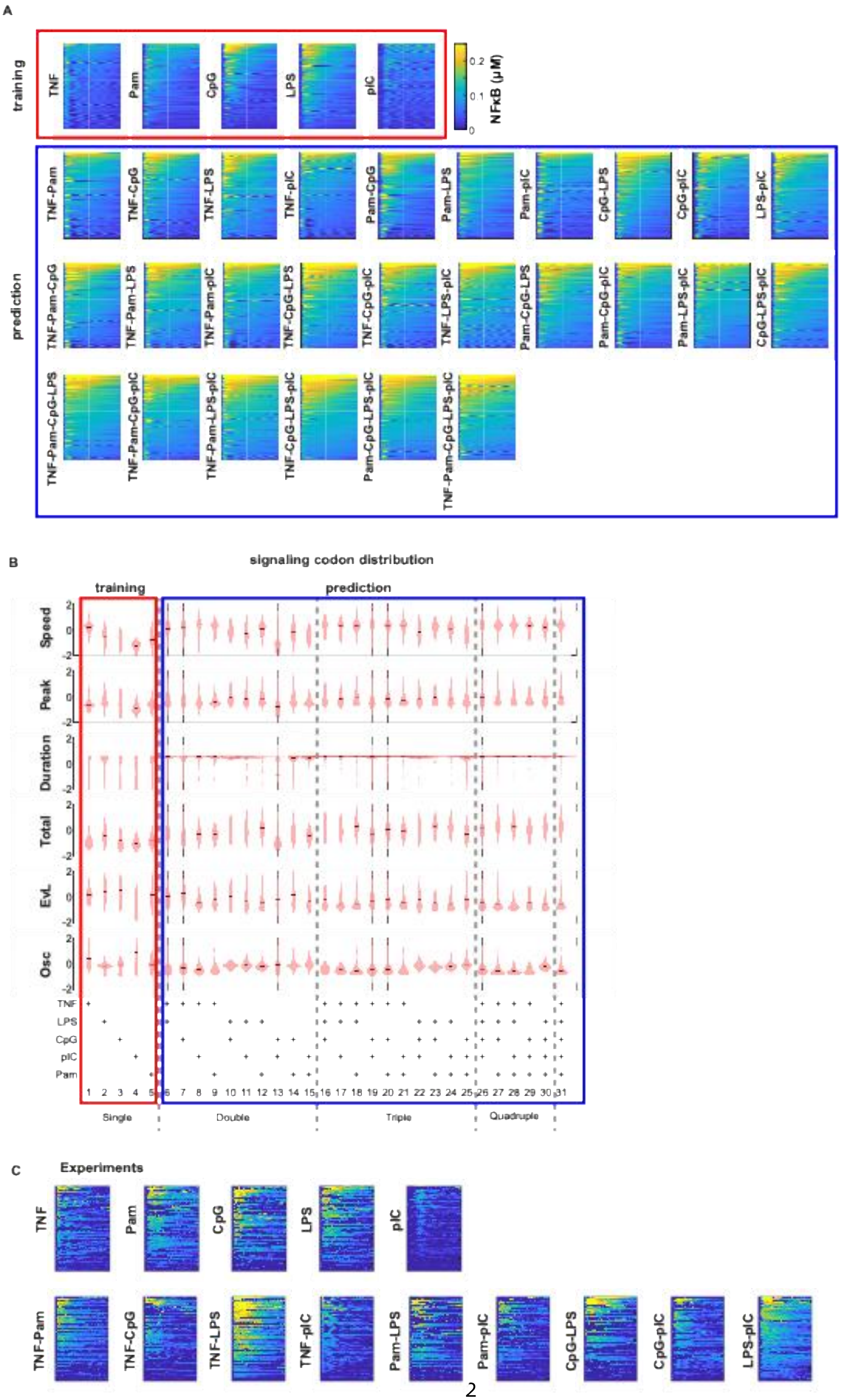
A
training
TNF
Pam
CpG
LPS
pIC
NFκB (μM)
0.2
0.1
0
prediction
TNF-Pam
TNF-CpG
TNF-LPS
TNF-pIC
Pam-CpG
Pam-LPS
Pam-pIC
CpG-LPS
CpG-pIC
LPS-pIC
TNF-Pam-CpG
TNF-Pam-LPS
TNF-Pam-pIC
TNF-CpG-LPS
TNF-CpG-pIC
TNF-LPS-pIC
Pam-CpG-LPS
Pam-CpG-pIC
Pam-LPS-pIC
CpG-LPS-pIC
TNF-Pam-CpG-LPS
TNF-Pam-CpG-pIC
TNF-Pam-LPS-pIC
TNF-CpG-LPS-pIC
Pam-CpG-LPS-pIC
TNF-Pam-CpG-LPS-pIC
B
signaling codon distribution
training
prediction
Speed
Peak
Duration
Total
EvL
Osc
2
0
-2
TNF
LPS
CpG
pIC
Pam
1 2 3 4 5 6 7 8 9 10 11 12 13 14 15 16 17 18 19 20 21 22 23 24 25 26 27 28 29 30 31
Single
Double
Triple
Quadruple
C
Experiments
TNF
Pam
CpG
LPS
pIC
TNF-Pam
TNF-CpG
TNF-LPS
TNF-pIC
Pam-LPS
Pam-pIC
CpG-LPS
CpG-pIC
LPS-pIC

#### Figure S1. Model prediction on combinatorial ligand stimulation

A. Heatmaps of simulated single-cell NFκB trajectories under single-ligand (training dataset, in red box) and combinatorial ligand (prediction dataset, in blue box) stimulation. For each heatmap, the condition is indicated on the left, and each row represents a single cell prediction, ordered by total activity of the signaling. In each subpanel, the x-axis represents time, and color intensity indicates NFκB abundance in accordance with the color bar.

B. Violin plots of signaling codon distributions from single ligand stimulation (training) to combinatorial ligand stimulation (prediction). Conditions are labeled at bottom. The mean value of the respective codon distribution is indicated.

C. Heatmaps of experimental single-cell NFκB trajectories under single-ligand (top row) and pairwise combinatorial ligand (bottom row) stimulation. Within each heatmap, each row represents a single cell prediction (ordered by total activity of the signaling); the x-axis represents time, and color intensity indicates NFκB abundance in accordance with the color bar in panel A.

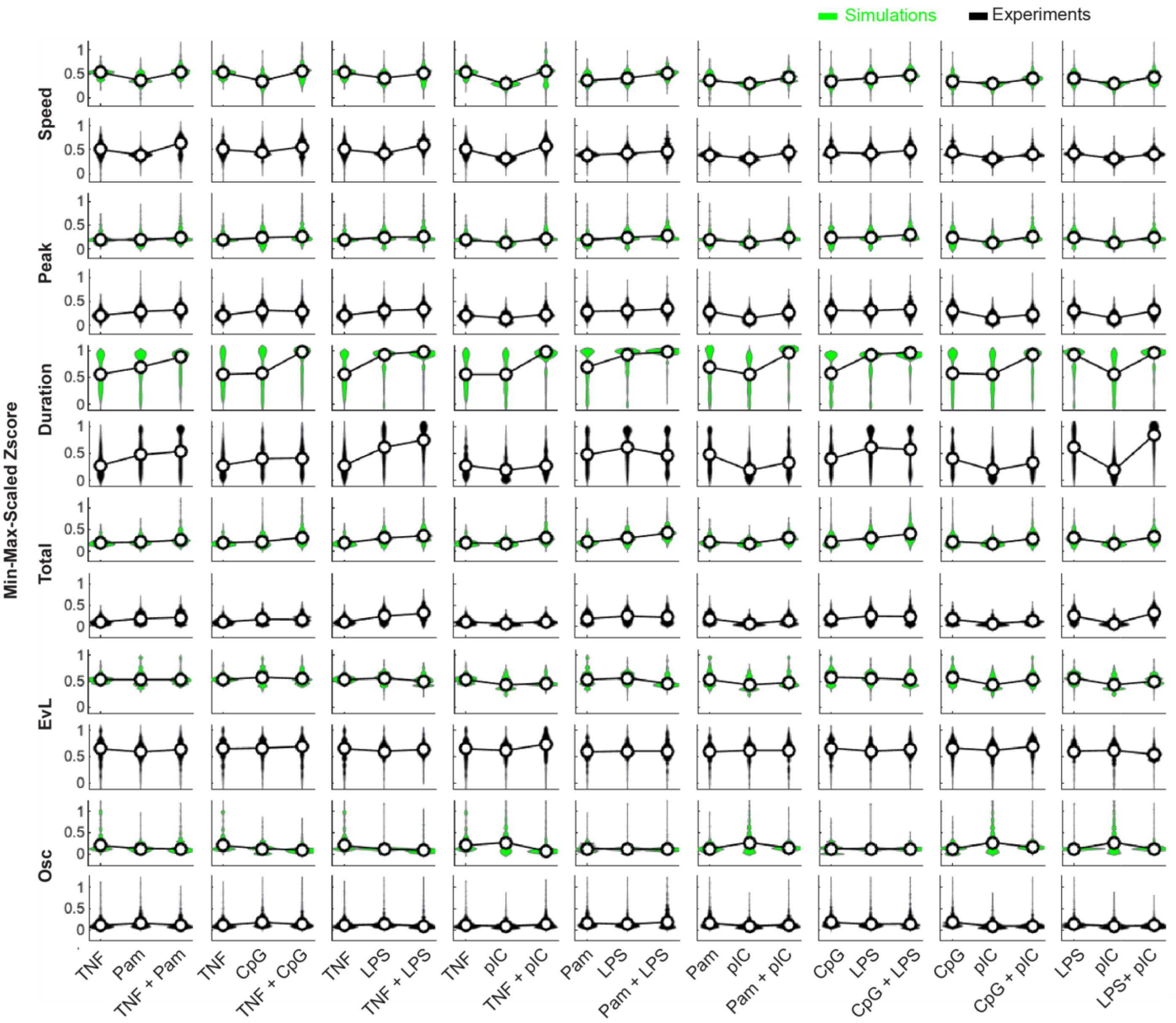

**Figure S2. Integrative and non-integrative NFκB signaling codons quantified from model simulations and experimental studies**

Violin plots of indicated signaling codon distributions from single and combinatorial ligand stimulation, as calculated from NFκB trajectories predicted by the model (green) or determined experimentally (black),. Each pair of rows corresponds to one signaling codon, detailed on the left. Columns correspond to different combinations. Within each subpanel, the first two columns represent single ligand stimulation, and the third column signifies dual ligand stimulation, with ligand details on the x-axis labels. The hollow circle in each plot symbolizes the median of the respective codon.

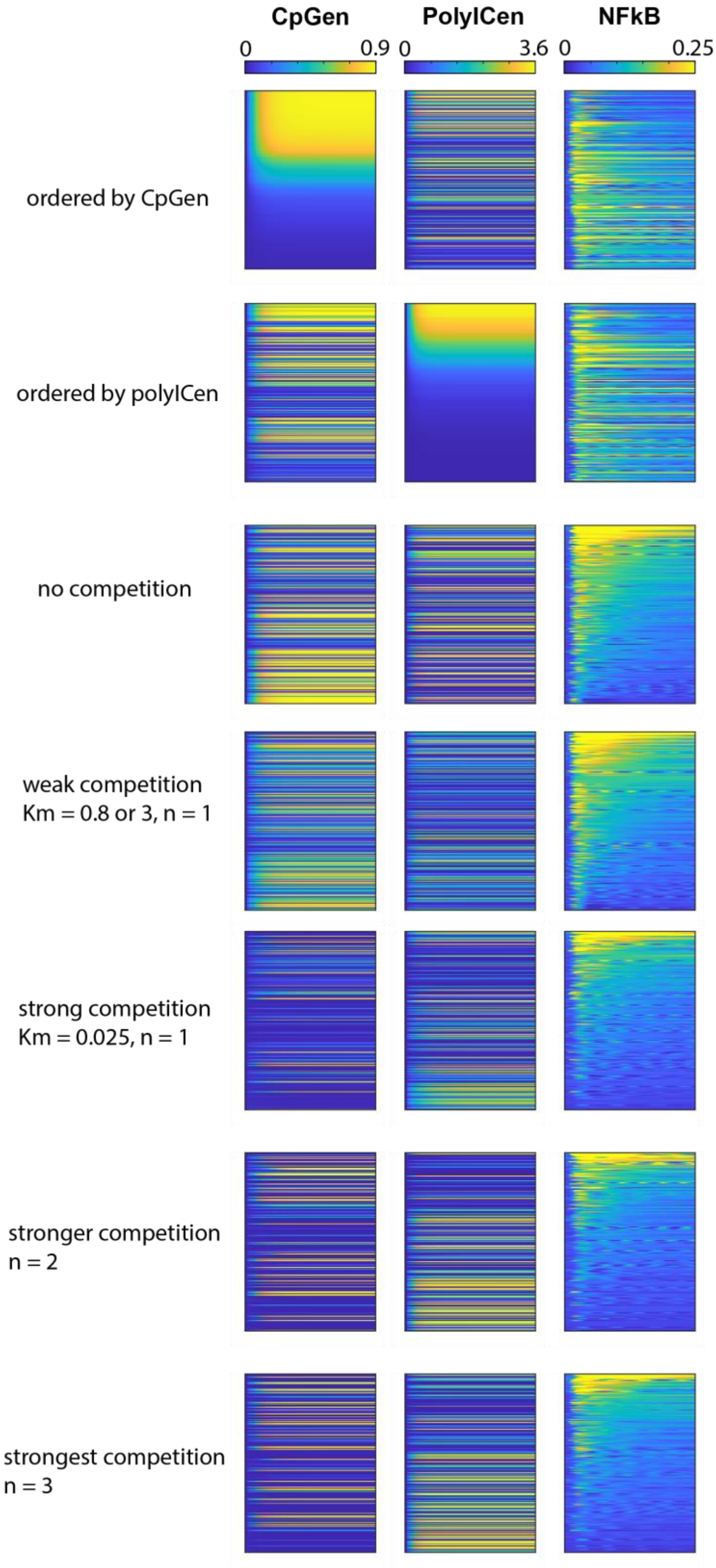

**Figure S3. Competition between CpG and pIC in endosomal transport**

Heatmaps of simulated single-cell endosomal CpG (left column), pIC activity (middle column), and NFκB trajectories (right column) under combined CpG-pIC stimulation. The first three rows correspond to the original model without competition between CpG and pIC, ordered by total integral of endosomal CpG activity (first row), total integral of endosomal PolyIC activity (second row), and total NFκB signaling (third row). Rows 4 to 7 are based on the updated model (Figure 2D) with varying the Hill coefficient, as indicated on the left. In each heatmap, rows correspond to single-cell predictions, the x-axis represents time, and color intensity indicates endosomal CpG, endosomal PolyIC, or NFκB abundance in accordance with the color bar specified on the top.

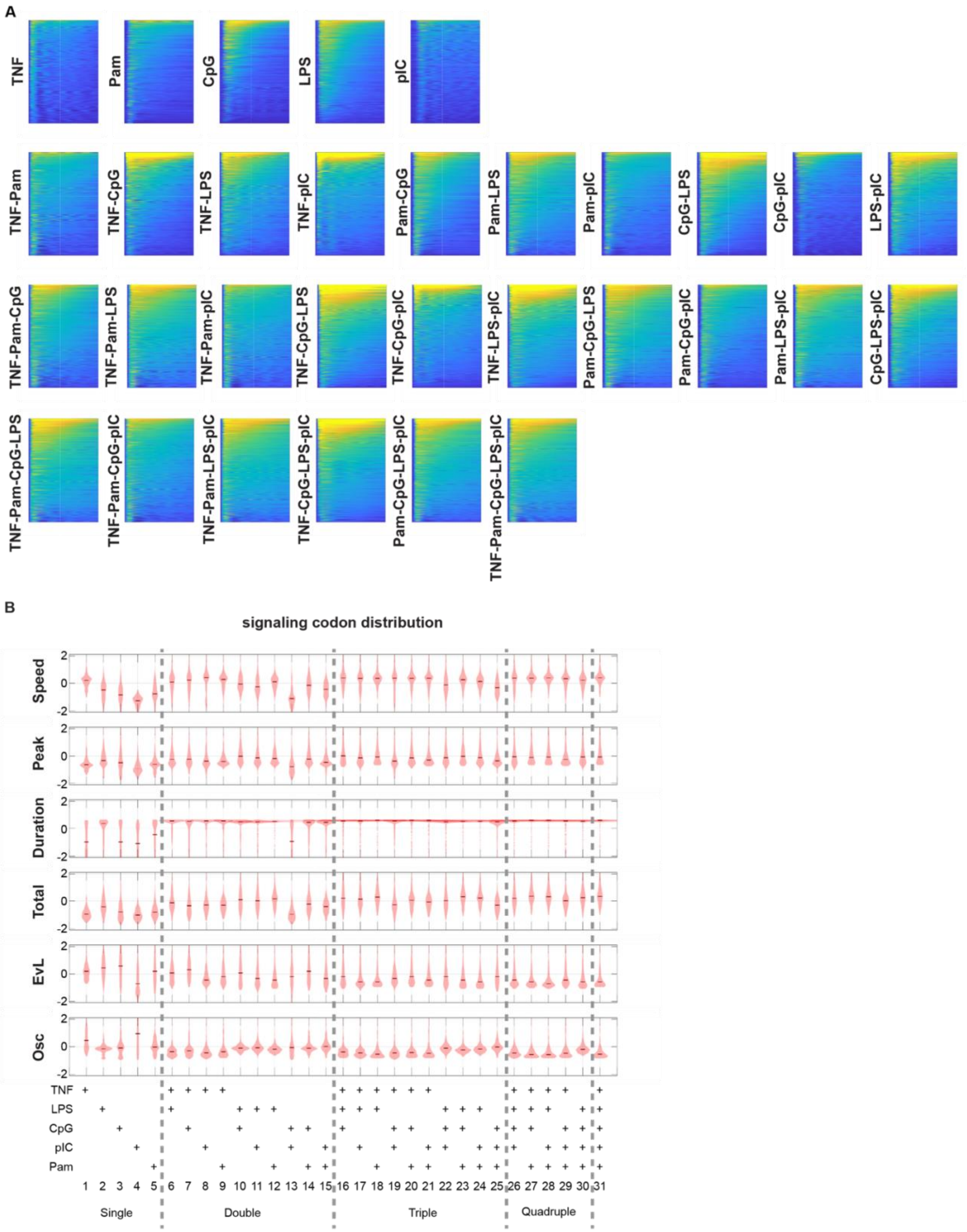

#### Figure S4. Updated model simulations of combinatorial ligand stimulations

A. Heatmaps of simulated single-cell NFκB trajectories under single and combinatorial ligand stimulation using the updated model (Figure 2D). For each heatmap, the condition is indicated on the left, and each row represents a single cell prediction, ordered by Total signaling activity. In each subpanel, the x-axis represents time, and color intensity indicates NFκB abundance in accordance with the color bar.

B. Violin plots of signaling codon distributions from signaling trajectories shown in panel A. Conditions are labeled at bottom. The mean value of the respective codon distribution is indicated.

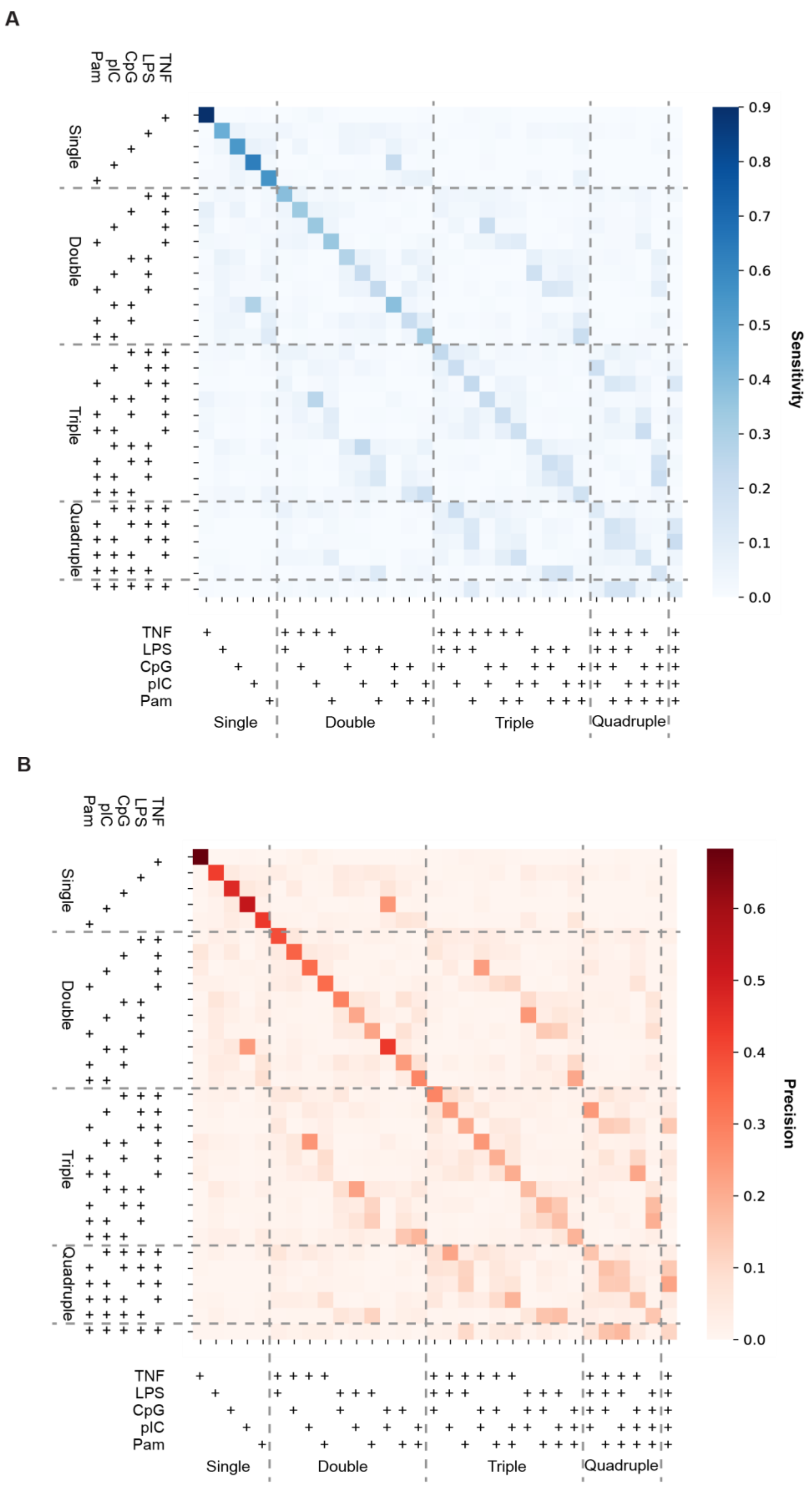

**Figure S5. Classifier predictions of combinatorial ligand responses**

A. Heatmap of Sensitivity matrix for the Random Forest classifier shown in Figure 3C, which was trained on all 31 conditions using NFκB response signaling codons as input to predict the stimuli.
B. Heatmap of Precision matrix for the same Random Forest classifier.

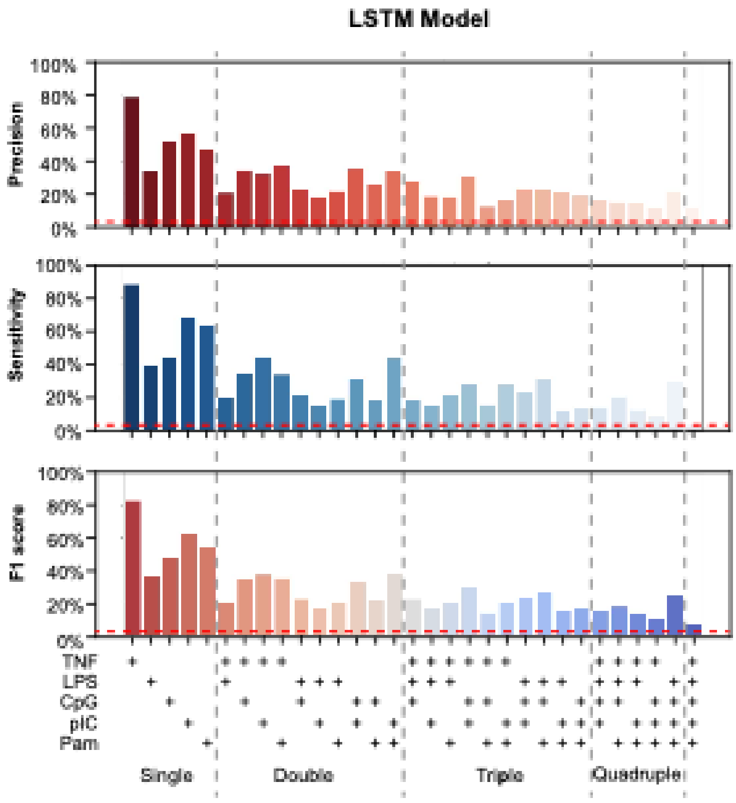

**Figure S6. Predicting stimuli using a Long Short-Term Memory (LSTM) model**

Bar plots of Precision, Sensitivity, and F1 score for Long Short-Term Memory (LSTM) model trained on all 31 conditions using NFκB time-series trajectories as input to predict the stimuli. 60% of the data are used for training and 40% for testing.

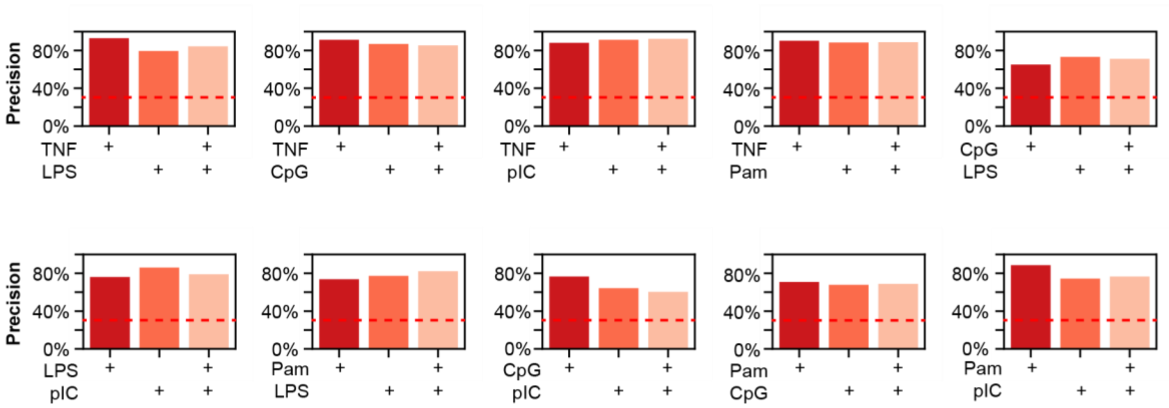

**Figure S7. Classifier predictions for dual-ligand stimulations with the corresponding single-ligand conditions**

Bar plots of precision for Random Forest classifier trained on all 3 conditions (different pairs of ligands, and their combination) using NFκB response signaling codons as input to predict the stimuli.

### Supplementary notes for materials and methods

#### Combinatorial ligand prediction

To generate heterogeneous single-cell NFκB trajectories, we first sampled signaling network parameters from an inferred distribution based on experimental data (1). These parameters included the core module and those from a single receptor module. For predicting responses to combinatorial ligand stimuli, additional receptor module parameters were assigned using equations (1–2). Once the heterogeneous virtual single-cell NFκB networks were established, we simulated their NFκB responses to ligand combinations.

To construct these virtual networks, we employed a permutation sampling approach, drawing inferred single-cell parameters from experimental data (1). Specifically, parameters were sampled from a non-parametric distribution with the density function:

$$f(\psi) = \sum_{m=1}^{M} \frac{1}{M} \delta\left(\psi - \hat{\psi}^{(m)}\right)$$

where $M$ represents the total number of cells across all dosage conditions for a given ligand. We used a permutation sampling strategy because simulations showed that it closely reproduced the experimental NFκB dynamic feature distributions, whereas sampling from a parameterized distribution led to substantial deviations (1).

Since experimentally inferred single-cell parameters included only the core module and one single receptor module, the missing receptor modules had to be inferred to reconstruct complete NFκB signaling networks. To generate $N$ heterogeneous single-cell NFκB networks for predicting responses to n-ligand stimuli, we first sampled $N/n$ parameter sets corresponding to the 1st receptor module and core module from the experimental data inferred parameter sets. These defined the core and one receptor module parameters for each virtual single cell. To infer parameters for the remaining $n-1$ receptor modules, we assigned values from 2nd ligand stimulated single cells with same—or the most similar—core module parameters (Equation 1-2 in main text). This process was repeated iteratively, treating each receptor module as the initial sampling module in turn, until N virtual single-cell NFκB signaling networks were fully reconstructed.

The sampled parameters were then applied to simulate single-cell NFκB signaling trajectories using a 52-dimensional ODE model (see Supplementary dataset 1). For each condition, between 999 and 1,000 cells were simulated ($N = 999$ for $n = 3$; $N = 1000$ for $n = 1,2,4,5$) using

Matlab's ode15s solver. Simulations included an initial phase to establish steady-state conditions, followed by stimulation. The resulting trajectories were visualized in MATLAB.

**Experimental data generation**

Macrophage Cell Culture and Stimulation

Myeloid precursor cells were prepared from the RelA-mVenus mouse strain (2) by HoxB4-mediated transduction (hMP) (3). hMP-Derived Macrophages (hMPDMs) were prepared by culturing hMPs in L929-conditioned medium using the standard Bone-Marrow Derived Macrophage (BMDM) culture method (2). hMPDMs were re-plated in imaging dishes on day 6 or day 7 at 12,000 to 18,000 cells/well in an 8-well ibidi SlideTek chamber, for imaging at an appropriate density on day 10, day 11, or day 12. hMPDMs were left untreated before stimulation. Stimulation was done with the toll-like receptor (TLR) 4 agonist, lipopolysaccharide (LPS) (10 ng/mL) (Sigma Aldrich), TLR3 agonist, polyinosine-polycytidylic acid (Poly(I:C)) (50 µg/mL) (InvivoGen), TLR9 agonist, CpG B ODN (100 nM) (InvivoGen); TLR2 agonists, Pam3CSK4 (100 ng/mL) (InvivoGen), or cytokine TNF (1 ng/mL) (Roche) without media replacement. Doses were selected to give maximal response.

Live-cell Imaging

Macrophages were stained with nuclear staining dye, Hoechst 33342 (5 ng/mL) two hours prior to imaging within ibidi chambers. Cells were imaged at 5-minute intervals on a Zeiss AxioObserver platform with live-cell incubation, using epifluorescent excitation from a Sutter Lambda XL light source. The first three images collected (pre-stimulation) were used to determine the baseline activity of NFκB for each cell. After 15 minutes of the start of imaging, conditioned culture media containing stimulus was injected into the respective well of the ibidi chamber in situ. Images were recorded on a Hamamatsu Orca Flash 2.0 CCD camera for 12.5 hours.

Image Processing

Microscopy time-lapse images were exported for single-cell tracking and measurement in MATLAB R2018a, used in earlier work (2). Briefly, cells were identified using DIC images, then segmented, guided by nuclear staining from the Hoechst image. Segmented cells were linked into trajectories across successive images, then nuclear and cytoplasmic boundaries were defined and used for measurement in the fluorescent channel for mVenus-NFκB. Nuclear NFκB levels were quantified on a per-cell basis, normalized to image background levels, then baseline-subtracted. The first three images collected (pre-stimulation) were used to determine the baseline activity of NFκB for each cell. The mean fluorescence value from these three frames was subtracted from the complete trajectory to normalize each cell. For downstream analysis and visualization, the third timepoint corresponds to time = 0 and 97 timepoints after that were included (8-hour trajectories).

Mitotic cells, as well as cells that drifted out of the field of view, were excluded from analysis. The code (MACKtrack) used for this analysis is publicly available at GitHub (https://github.com/brookstaylorjr/MACKtrack ).

**Signaling codon decomposition and Responder fraction**

Simulation responder ratio is calculated via peak value or integral higher than specific threshold, implemented in MATLAB. Experimental data are first rescale to the similar numerical range under standard units (4, 5, 1) then calculated responder ratio. Six NFκB signaling codons—Speed, Peak, Duration, Total, EvL, and Osc—encoding stimulus information (2) are calculated. The calculation is the same as (1), and detailed in the codes (https://github.com/Xiaolu-Guo/Combinatorial_ligand_NFkB). Specifically, 'duration' measures the total time of NFκB level over a low threshold defining Duration; 'time2HalfMaxPosInt' measures how much NFκB activity is "front-loaded", defining EvL; 'oscpower' distinguishes oscillation features of trajectory, defining Osc; 'max-value' and 'pos-pk1-amp' measures the peak amplitude, defining Peak; 'max-pos-pk1-speed', 'pos-pk1-time', 'derivatives', measures the activation speed, defining Speed; 'max-pos-integral' measures the accumulated activity, defining Total.

**Competition between CpG and pIC**

We use the Hill equation formula to model the competition between CpG and pIC due to endosomal capacity saturation. Specifically, CpG imported to the endosome are assumed to reduce the cell's capacity for pIC import to endosome, and vice versa. Thus, we multiply the endosomal transportation rate by the inhibition Hill formula:

$$\frac{d(CpG_{en})}{dt} = f_{CpG_{en}}(CpG_{en}, TLR9, TLR9_{CpG}) + f_{CpG,in}(CpG) * \left(\frac{v_{pIC}}{K_{pIC}^{n_{pIC}} + pIC_{en}^{n_{pIC}}}\right)$$

$$\frac{d\,(pIC_{en})}{dt} = f_{pIC_{en}}(pIC_{en}, TLR3, TLR3_{pIC}) + f_{pIC,in}(pIC) * \left(\frac{v\_CpG}{K_{CpG}^{n_{CpG}} + CpG_{en}^{n_{CpG}}}\right)$$

The full formula of these two equations are:

$$\frac{d(CpG_{en})}{dt} = -\psi_{89,1} * CpG_{en} - \psi_{90,1} * CpG_{en} * TLR9 + \frac{\psi_{88,1} * CpG^{\psi_{88,3}}}{CpG^{\psi_{88,3}} + \psi_{88,2}^{\psi_{88,3}}} * \left(\frac{\psi_{88,5}}{\psi_{88,7}^{\psi_{88,6}} + polyIC_{en}^{\psi_{88,6}}}\right) + \psi_{91,1} * TLR9_{CpG}$$

and

$$\frac{d\,(polyIC_{en})}{dt} = -\psi_{80,1} * polyIC_{en} - \psi_{81,1} * polyIC_{en} * TLR3$$

$$+\frac{\left(\psi_{79,1} * polyIC^{\psi_{79,3}}\right)}{polyIC^{\psi_{79,3}} + \psi_{79,2}^{\psi_{79,3}}} * \left(\frac{\psi_{79,5}}{\psi_{79,7}^{\psi_{79,6}} + CpG_{en}^{\psi_{79,6}}}\right)$$
$$+ \psi_{82,1} * TLR3_{polyIC}$$

These equations replaced the original ODE equations of $\frac{d(CpG_{en})}{dt}$ and $\frac{d\,(polyIC_{en})}{dt}$ for the updated model.

**Stimulus-response specificity assessment**

To analyze stimulus-response specificity of NFκB dynamics across different stimulations, we computed the distributions of signaling codons from simulated datasets using the updated CpG-pIC competition model. Wasserstein-2 distances were computed between codon distributions across different conditions, and the average distance across six signaling codons was used to quantify stimulus-response specificity.

A random forest classifier was trained on simulated data to predict conditions with NFκB signaling codons as input. The model was implemented using 'RandomForestClassifier' in python. The dataset included 31 ligand mixture conditions: five single ligands—TNF, LPS, CpG, PolyIC, and Pam3CSK—and 26 ligand combinations of these single ligands. The dataset was split into 67% training and 33% testing, comprising 30,990 virtual cells, with 999~1,000 vells per condition.

Model performance was assessed using a confusion matrix, where predicted vs. true ligand classification accuracy was recorded. Evaluation metrics included sensitivity, precision, and F1-score (harmonic mean of precision and sensitivity) to measure per-class performance. Heatmaps were used to visualize classification trends and misclassification patterns.

For pair-ligand combinations and the corresponding single-ligand condition, the same classification approach was applied to the corresponding three condition -- two single ligands and their combination -- for each pair. This resulted in a total of 10 unique base-paired ligand combinations.

For classifier on predicting specific ligand presence or not in ligand mixtures, we grouped all double-ligand, triple-ligand, and quadruple-ligand conditions (in total 25 conditions) into two categories: one with 14 conditions where the specific ligand is present and another with 11 conditions where that ligand is absent. We then trained machine learning classifiers to predict whether a cell was exposed to that ligand based on its NFκB signaling codon. 5 different classifiers were trained for 5 ligands.

Similarly, LSTM classifiers were used to classify conditions based on NFκB dynamic trajectories. This was implemented using 'LSTM' in 'tensorflow.keras.layers' in python. The dataset was split into 60% training and 20% validation and 20% testing. All model performance metric are calculated using the 20% validation + 20% testing, in total 40% of the dataset.
